# Estimation of biodiversity metrics by environmental DNA metabarcoding compared with visual and capture surveys of river fish communities

**DOI:** 10.1101/617670

**Authors:** Hideyuki Doi, Ryutei Inui, Shunsuke Matsuoka, Yoshihisa Akamatsu, Masuji Goto, Takanori Kono

**Author notes:** Corresponding authors: Hideyuki Doi, Ryutei Inui. These authors equally contributed to this study (Hideyuki Doi, Ryutei Inui, Shunsuke Matsuoka).

## Abstract

1. Information on alpha (local), beta (between habitats), and gamma (regional) diversity is fundamental to understanding biodiversity as well as the function and stability of community dynamics. Methods like environmental DNA (eDNA) metabarcoding are currently considered useful to investigate biodiversity.
2. We compared the performance of eDNA metabarcoding with visual and capture surveys for estimating alpha and gamma diversity of river fish communities, and nestedness and turnover in particular.
3. In five rivers across west Japan, by comparison to visual/capture surveys, eDNA metabarcoding detected more species in the study sites (i.e., alpha diversity). Consequently the overall number of species in the region (i.e., gamma diversity) was higher. In particular, the species found by visual/capture surveys were encompassed by those detected by eDNA metabarcoding.
4. Estimates of community diversity within rivers differed between survey methods. Although we found that the methods show similar levels of community nestedness and turnover within the rivers, visual/capture surveys showed more distinct community differences from upstream to downstream. Our results suggest that eDNA metabarcoding may be a suitable method for community assemblage analysis, especially for understanding regional community patterns, for fish monitoring in rivers.

## Introduction

The maintenance of biodiversity underpins the stability of ecosystem processes in constantly changing environments (Primack, 1993; Margules & Pressey, 2000; Pecl et al., 2017). Moreover, biodiversity loss affects ecosystem functions and services and, consequently, human society (Primack 1993; Margules & Pressey, 2000, Pecl et al. 2017). Ecologists have made efforts to conserve biodiversity based on essential biodiversity survey methods, especially for species richness and distribution (Primack, 1993; Margules & Pressey, 2000, Doi & Takahara, 2016, Pecl et al., 2017). Biodiversity can be evaluated in different levels: e.g., by estimating alpha (local), beta (between habitats), and gamma (regional) diversity and the variation of the community assemblages. To conserve local communities, ecologists incorporate these diversity metrics into management decisions (Primack 1993; Margules & Pressey, 2000, Socolar et al., 2016). For example, measuring variation among community assemblages can identify biodiversity loss and inform the placement of protected areas and the management of biological invasions and landscapes (Socolar et al., 2016). Thus, robust methods for monitoring biodiversity are fundamental for biodiversity and environmental management.

Environmental DNA (eDNA) analysis is considered a useful tool to investigate the distribution and richness of aquatic and terrestrial organisms (Takahara et al., 2012, 2013; Rees et al., 2014; Goldberg et al., 2015; Miya et al., 2015; Thomsen & Willerslev, 2015; Doi et al., 2017; Doi et al., 2019; Fujii et al., 2019). High-throughput sequencing derived from eDNA, called “eDNA metabarcoding”, is an exceptionally useful and powerful tool for community biodiversity surveys (Taberlet et al., 2012; Deiner et al., 2016, 2017; Sato et al., 2017; Bylemans et al., 2018; Fujii et al., 2019). eDNA metabarcoding has recently been applied in fish community surveys, e.g. Miya et al. (2015) designed and applied universal PCR primers (the MiFish primers) to survey marine fish communities. To confirm the usefulness of eDNA metabarcoding for community assessment, many studies have compared eDNA metabarcoding to a species list generated by traditional surveys including visual and capture methods (Deiner et al., 2016; Bylemans et al., 2018; Nakagawa et al., 2018; Yamamoto et al., 2018; Fujii et al., 2019). However, comparisons of eDNA and traditional survey methods for estimating regional (gamma) diversity or differences in community assemblage composition among locations are more limited (but see Drummond et al., 2015; Deiner et al., 2016; Staehr et al. 2016, Maechler et al. 2019).

Variation in community assemblage composition reflects two components, nestedness and species turnover for considering regional structure of community assemblage (Harrison et al., 1992; Baselga et al., 2007; Baselga, 2010). Nestedness occurs when the community at the sites with fewer species are subsets of the community at the sites with higher species richness (Wright & Reeves, 1992; Ulrich & Gotelli, 2007) and generally reflects a non-random process of species loss (Gaston & Blackburn, 2000). Contrastingly, species turnover implies the replacement of some species by others because of environmental sorting or spatial/historical constraints (Baselga, 2010). Statistical separation methods for nestedness and species turnover were applied for evaluating variation of the community assemblages in various systems (Baselga, 2010; Baselga et al., 2012). Moreover, Baselga’s (2010) framework can be applied to compare the performance among methods when evaluating beta diversity via nestedness and species turnover. Nestedness and species turnover can also occur due to differences in sampling methods, even when the surveys are conducted on the same community. For example, differing biases in species detection between two methods can result in apparent species turnover. Evaluating nestedness and species turnover would allow us to reveal the potential bias of community analysis of eDNA metabarcoding compared to traditional surveys. Therefore, we can quantitatively compare the performance of eDNA metabarcoding and traditional surveys for alpha/gamma diversity evaluation of biological communities and the variation of the community assemblages among the study sites.

Using statistical methods, we tested the performance of eDNA metabarcoding in five river systems in different regions with various fish species. We conducted eDNA metabarcoding using universal MiFish primers that target fish and identified the fish by visual snorkeling and hand-net capture surveys. We evaluated the performance of eDNA metabarcoding by comparing the obtained fish community structure to that evaluated by visual/capture survey with special regard to nestedness and species turnover.

## Methods

### Site description

In 2016, we conducted field surveys in five river systems across Japan (river map in Fig. S1): the Kyuragi River on October 10, the Koishiwara River on October 21, the Yato River on October 25, the Hazuki River on November 2, and the Oze River on November 6. The survey sites were set at a site at each of three river segments (upstream, mid-stream, and downstream, the internal distances ranged from 4.5 to 25.8 km, Fig. S1) for each river. Each site was set so that the length in the up-down direction was approximately 100 m with a riffle at the downstream end (e.g., Fig. S2).

### Water collection for eDNA survey

In each site, we collected 1 L of surface water in bleached bottles at two points, in flowing water near the downstream end of the site, and in stagnant or nearly stagnant water near the shore (Fig. S2) immediately before visual and capture surveys. Prior to sample collection, eDNA was removed from the bottles and filtering equipment using 10% commercial bleach (ca. 0.6% hypochlorus acid) and washing with DNA-free distilled water. One milliliter of benzalkonium chloride (BAC, 10% w/v) was added per liter of water sample to avoid a decrease in eDNA concentration in the samples (Yamanaka et al., 2016). During transport, samples were stored in a cooler with ice packs. The ‘cooler blank’ contained 1 L DNA-free water, which we brought to the field and treated identically to the other water samples, except that it was not opened at the field sites.

### Visual observation and capture methods

After water sampling, the fish fauna survey was conducted by visual observation with snorkel and collection with hand net. For visual observation, we observed and recorded fish species by snorkeling in a 100-m transect (snorkeling by 1 person for 1 h, Fig. S2). We observed at various micro habitats, including the riffle, pool, and shore bank from the downstream end to upstream end. We also conducted a hand-net capture survey (1 person × 1 h) using a D-frame net (2 mm mesh, net opening: 0.16 m^2^) in the various habitats in the river, including the riffle, pool, and shore bank. Fishes were identified according to Nakabo et al. (2013) at the survey site. We used the combined taxa list from both traditional surveys to compare to that of eDNA metabarcoding. In order to prevent contamination of eDNA samples, the investigator who collected and identified the fish (i.e., handled the fish) and the investigator who sampled the water were different.

### eDNA collection, extraction and measurements

Collected water samples were vacuum-filtered into GF/F glass filters (47 mm diameter, pore size: 0.7 μm, GE Healthcare, Little Chalfont, UK) in the laboratory within 24 h of sampling. After filtration, all filters were stored at −20 °C before eDNA extraction. The cooler blank was also processed in the same manner. A liter of Milli-Q water was used as the filtering blank to monitor contamination during filtering in each site and during subsequent DNA extraction.

To extract the DNA from the filters, we followed the methods described in Uchii, Doi, & Minamoto (2016). We incubated the filter by submerging the mixed buffer of 400 μL of Buffer AL in DNeasy Blood & Tissue Kit (Qiagen, Hilden, Germany) and 40 μL of Proteinase K (Qiagen, Hilden, Germany), using a Salivette tube (Sarstedt, Nümbrecht, Germany) at 56 °C for 30 min. The Salivette tube with filters was centrifuged at 5000 × *g* for 5 min. Then, we added 220 μL of TE buffer (pH: 8.0) onto the filter and again centrifuged at 5000 × *g* for 5 min. The DNA was purified using a DNeasy Blood & Tissue Kit with extracted the DNA in 200 μL in Buffer AE. Samples were stored at −20 °C until the 1st-PCR assay.

### Library preparation and MiSeq sequencing

The detailed molecular methods are described in Fujii et al. (2019) with a two-step PCR-procedure for Illumina MiSeq sequencing. Briefly, we performed 1st-PCR withMiFish-U-F and MiFish-U-R primers (Miya et al., 2015), which were designed to contain Illumina sequencing primer regions and 6-mer Ns;

Forward: 5′-*ACACTCTTTCCCTACACGACGCTCTTCCGATCT* NNNNNN GTCGGTAAAACTCGTGCCAGC-3′,

Reverse: 5′-*GTGACTGGAGTTCAGACGTGTGCTCTTCCGATCT* NNNNNN CATAGTGGGGTATCTAATCCCAGTTTG-3′

The italicized and non-italicized letters represent MiSeq sequencing primers and MiFish primers, respectively, and the six random bases (N) were used to enhance cluster separation on the flow cells during initial base call calibrations on the MiSeq (Miya et al. 2015, Doi et al. 2019).

We performed the 1st-PCR with a 12 μL reaction volume containing 1× PCR Buffer for KOD FX Neo (Toyobo, Osaka, Japan), 0.4 mM dNTP mix, 0.24 U KOD FX Neo polymerase, 0.3 μM of each primer, and 2 μL template. The thermocycling conditions for this step were as follows: initial denaturation at 94 °C for 2 min, followed by 35 cycles of denaturation at 98 °C for 10 s, annealing at 65 °C for 30 s, and elongation at 68 °C for 30 s, followed by final elongation at 68 °C for 5 min. The first PCRs were performed using eight replicates (Doi et al. 2019) and individual first PCR replicates were pooled and purified using AMPure XP (Beckman Coulter, Brea CA, USA) as templates for the 2nd-PCR. The Illumina sequencing adaptors and the eight bp identifier indices (XXXXXXXX) were added to the subsequent PCR process using a forward and reverse fusion primer:

Forward: 5′-*AATGATACGGCGACCACCGAGATCTACA* XXXXXXXX ACACTCTTTCCCTACACGACGCTCTTCCGATCT-3′

Reverse: 5′-*CAAGCAGAAGACGGCATACGAGAT* XXXXXXXX GTGACTGGAGTTCAGACGTGTGCTCTTCCGATCT-3′

The italicized and non-italicized letters represent MiSeq P5/P7 adapter and sequencing primers, respectively. The 8X bases represent dual-index sequences inserted to identify different samples (Hamady et al. 2008). We performed the 2nd-PCR with 12 cycles of a 12 μL reaction volume containing 1× KAPA HiFi HotStart ReadyMix, 0.3 μM of each primer, and 1.0 μL of the first PCR production. The thermocycling conditions profile after an initial 3 min denaturation at 95 °C was as follows: denaturation at 98 °C for 20 s, annealing, and extension combined at 72 °C (shuttle PCR) for 15 s, with the final extension at the same temperature for 5 min. We confirmed the positive bands of the targeted 1st-PCR amplicons by electrophoresis. The 2nd-PCR products were pooled in equal volumes and purified using AMPure XP.

The purified PCR products were loaded on a 2% E-Gel SizeSelect (Thermo Fisher Scientific, Waltham, MA, USA) and the target size of the amplified DNA (approximately 370 bp) was collected. The samples concentration and quality were estimated by a Qubit dsDNA HS assay kit and a Qubit 2.0 (Thermo Fisher Scientific). The amplicons were sequenced by 2 × 250 bp paired-end sequencing on the MiSeq platform using the MiSeq v2 Reagent Kit. Note that the sequencing run contained a total of 339 samples including 40 of our samples (30 samples plus five cooler and five filter negative controls) and 299 samples from other research projects. The MiSeq sequencing was conducted in the Department of Environmental Solution Technology, Faculty of Science and Technology, Ryukoku University. All sequence data were deposited in DNA Data Bank of Japan (DRA, Accession number: DRA008090).

### Bioinformatic analysis for MiSeq sequencing

The detailed procedures used for bioinformatics analysis are described in Fujii et al. (2019). First, low-quality tails were trimmed from each read and paired-end reads were then merged. For the obtained 1,823,446 reads, primer sequences were removed and identical sequences (i.e., 100% sequence similarity) were merged using UCLUST (usearch 7.0.1001, Edgar, 2010). The sequences with 10 or more identical reads were subjected to the downstream processes. To annotate the taxonomy, local BLASTN search using BLAST 2.2.29 was conducted with the reference database of fish species for processed reads (Miya et al., 2015). The top BLAST hit with a sequence identity ≥ 97% was applied to species detection of each sequence. Note that the species were mostly identified with a ≥ 99% match. From the BLAST results, we identified the species using methods previously described (Sato et al., 2017). Also, we detect most of the fish species known to occur in the rivers of the region, according to Kawanabe et al. (2001).

### Statistical analyses

All statistical analyses and graphics were conducted in R ver. 3.4.4 (R Core Team, 2018). All statistics were set at the significance level α = 0.05. To compare between eDNA metabarcoding and visual survey data, the taxonomic levels in the species list from visual survey were compared to the lists from eDNA metabarcoding (Table S1, S2) in reference to previous studies using the MiFish primer (Sato et al., 2017; Fujii et al., 2019). Before statistical analysis, we confirmed that the sequencing depth was sufficient to detect alpha diversity in the samples by “iNEXT” and “ggiNEXT” functions in the “iNEXT” ver. 2.0.19 package (Chao et al. 2014, Fig. S3). We merged the community data from two points, the stream near the downstream end and the shore, to compare with visual/capture surveys.

We tested the differences in fish richness of sites, segments, and rivers between both methods using generalized linear mixed models (GLMMs) with the “lmer” function in the “lme4” ver. 1.1-21 package (Bates et al. 2015). In the GLMM models, the method was treated as a fixed effect with Poisson distribution, and the rivers and segments were treated as random effects.

The differences in community compositions were visualized using nonmetric multidimensional scaling (NMDS) with 500 separate runs of real data. For NMDS, the two data sets (eDNA metabarcoding and visual/capture survey) were combined into one ordination (but showed divided into two panels by “facet_grid” function in ggplot2). The community dissimilarity was calculated based on incidence-based Jaccard indices. We evaluated the differences in community structures between methods and sites using permutational multivariate analysis of variance (PERMANOVA). For PERMANOVA, we used Jaccard and Raup-Crick dissimilarity indices and calculated the statistical values with 999 permutations. The Raup-Crick index estimates the probability that two samples have greater differences in species composition than would be expected by chance draws from the regional species pool, and accounts for differences in alpha diversity between samples. We used “metaMDS” and “adonis” functions in the “vegan” ver. 2.5-6 package (https://github.com/vegandevs/vegan) for NMDS ordination and PERMANOVA, respectively.

Indicator taxa analysis (Cáceres & Legendre, 2009) was performed to determine which taxa had significantly different frequencies between eDNA and visual/capture methods. The analysis was performed using the “signassoc” function in the “indicspecies” ver. 1.7.8 package on the presence/absence data for the testing with regarding the package description and Cáceres & Legendre (2009). The “signassoc” function can calculate the index with both presence/absence and abundance data. We used the presence/absence data with mode = 1 (group-based) and calculated the *P*-values with 999 permutations after Sidak’s correction of the multiple testing.

To compare community nestedness and turnover (Baselga, 2010; Baselga, 2012) as estimated by eDNA and visual/capture surveys, we used the “beta.pair” function in the “betapart” ver. 1.5.1 package (Baselga and Orme 2012). Standardized effect sizes (SESs) were calculated to show the degree of nestedness and turnover structure. The significance was defined by deviation from zero, and the expectation of random assembly (a null model) was estimated with 999 random sampling replicates. The SES was defined as follows: (β_obs_ − β_null_)/β_sd_, where β_obs_ is the observed beta diversity (here, the variation of the community assemblages among the sites), β_null_ is the mean of the null distribution of beta diversity, and β_sd_ is the standard deviation of the null distribution. SES values greater than zero indicate statistically stronger nestedness or turnover structure than expected under a random model of community assembly, while negative values indicate weaker nestedness or turnover than expected. The randomized community data were generated with independent swap algorithm (Gotelli, 2000) using “randomizeMatrix” function in the “picante” ver. 1.8 package (Kembel et al. 2010). First, to evaluate the differences in nestedness and species turnover between the survey methods (eDNA metabarcoding vs. visual/capture survey) at the same segments, the SES of pairwise nestedness and turnover were calculated for each sample pair within the river. Then, the fish community longitudinal nestedness and turnover structure along with river flow (i.e., upstream to downstream) by each method was evaluated with NODF and pairwise indices of nestedness. First, a nestedness metric (NODF, Almeida-Neto et al., 2008) and their SES value were calculated with 999 randomizations using the “nestednodf” and the “oecosimu” function in the “vegan” package. Then, we tested the differences in SES of pairwise indices between the survey methods (eDNA metabarcoding vs. visual/capture survey) by generalized linear mixed model (GLMM, with Gaussian distribution) with the “lmer” function in the “lme4” package. In the GLMM models for the SES of pairwise indices, the survey method was treated as a fixed effect, and the rivers-pairs and segment-pairs (i.e., three pairs in each river) were treated as random effects.

## Results

### Overview

We detected 53 fish taxa, almost all identified to the species or genus level, by eDNA metabarcoding in five rivers (Table S1, and S2) and visually observed 38 fish taxa in total. MiSeq paired-end sequencing for the library (30 samples plus five cooler and five filter negative controls) yielded a total of 1,601,816 reads (53,351 ± 17,639; mean ± S. D. for each sample, Table S2). We confirmed very low reads from negative controls (Table S2) with only one detection of a fish species, *Tridentiger* sp. in the blank of the Yato River, probably because of the cross-contamination among the samples. The reads of *Tridentiger* sp. in water samples (4783-9818 reads) were higher than in the blank sample (1286 reads), and so we retained this species in the analysis.

### Diversity indices between methods

We consistently detected more taxa at a site (i.e., higher alpha diversity) with eDNA metabarcoding than with visual/capture surveys (Fig. 1; GLMM, t = –5.45, *P* = 0.000018). Estimated taxa richness generally increased from upstream to downstream (Fig. 1, t = –5.85, *P* = 0.000004, random effect for variation among segments: 26.879 ± 5.185, SD). The random effect for “river” was not significantly different from zero (t = 1.737, *P* = 0.0942, random effect for variation among river: 3.896 ± 1.974).

**Figure 1.**
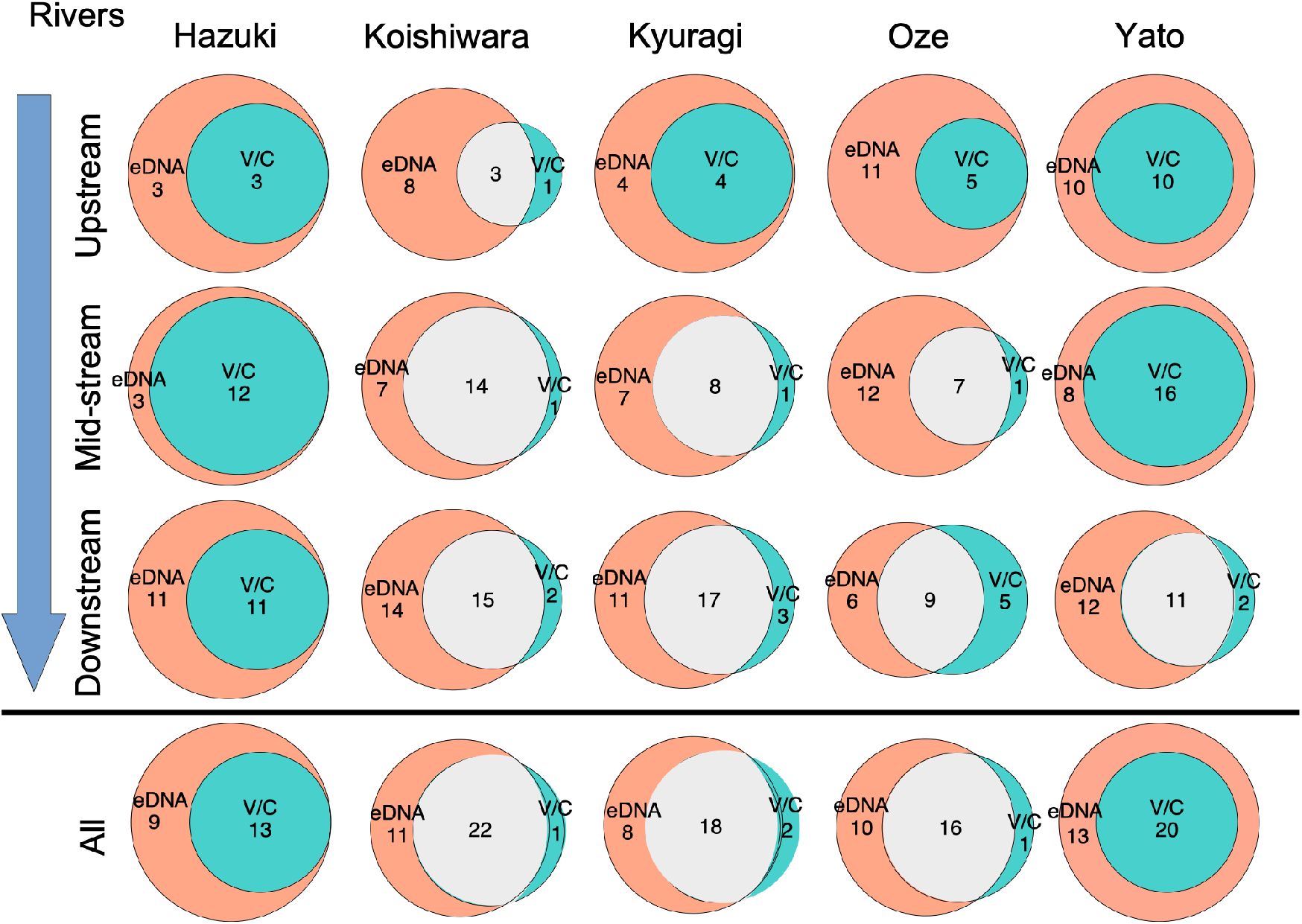
Numbers of taxa detected by eDNA metabarcoding (red), visual/capture surveys (blue), or both methods (white) at individual sites and for sites combined (ALL) within each of five river systems in Japan. Total taxa detected by eDNA metabarcoding is the sum of values included within the larger circle. Total taxa detected by visual/capture surveys is the sum of values in blue and white areas.

We ordinated the differences in community structure between the two methods by NMDS ordination by Jaccard index (Fig. 2) as well as Raup-Crick (Fig. S4). The patterns of differences in the both Jaccard and Raup-Crick indices were similar. The ordination showed the differences in community structure between the two methods, probably due to the differences in species detection in the communities. The PERMANOVA results with Jaccard and Raup-Crick indices for the ordination suggested there were differences in community composition evaluated by each method, eDNA metabarcoding and visual/capture survey (*P* < 0.012, Table S3). Moreover, communities from the combined results of eDNA metabarcoding and visual/capture survey were significantly different among rivers and segments (*P* < 0.029, Table S3). We found different patterns in ordinated river sites for each method (Fig. 2 for Jaccard index, Fig. S4 for Raup-Crick index).

**Figure 2.**
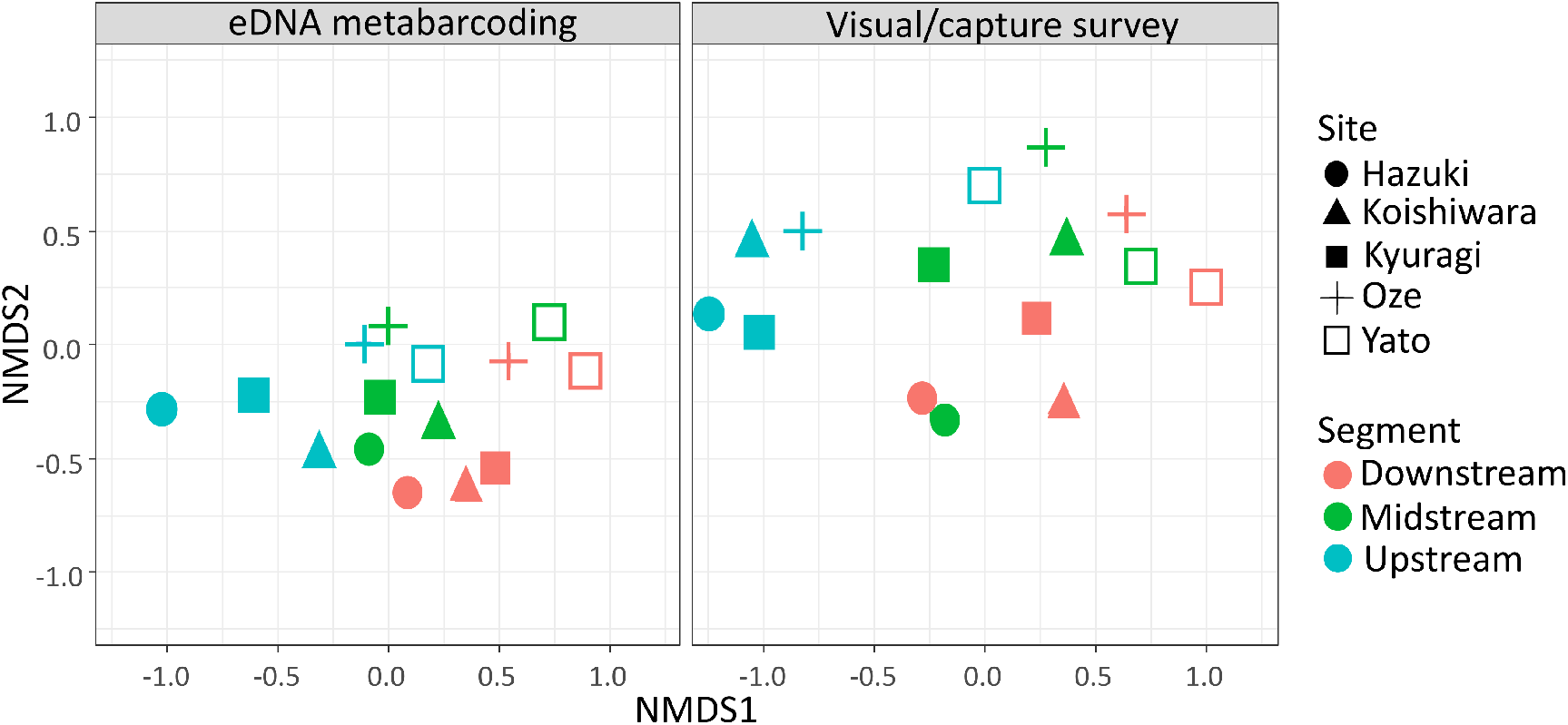
Nonmetric multidimensional scaling (NMDS) ordination of fish communities evaluated by study river (shapes) and segment (colors), using Jaccard index (ordination with Raup-Crick index shown in Fig. S4). A single ordination is divided into two panels to display differences in communities as measured by sampling methods. MDS stress was 0.158.

The PERMANOVA results with Jaccard index determined that communities were significantly different among the five rivers by eDNA metabarcoding (*P* = 0.001) but not by visual/capture survey (*P* = 0.12, Table S4). Conversely, the communities were significantly different among river segments (from downstream to upstream) by visual/capture survey (*P* = 0.011) but not significantly different (albeit marginally) by eDNA metabarcoding (*P* = 0.061, Table S4). The differences in PERMANOVA results with Jaccard index suggested that differences in the variation of the community assemblages among rivers across regions can be detected by eDNA metabarcoding but not by visual/capture survey. While, the PERMANOVA results with Raup-Crick index determined that communities were significantly different among the rivers by both eDNA metabarcoding (*P* = 0.001) and visual/capture survey (*P* = 0.018, Table S4).

Indicator taxa analysis comparing the communities estimated by both methods showed that eDNA metabarcoding more frequently detected several species than visual/capture survey, including Japanese eel (*Anguilla japonica*), salmon (e.g., *Oncorhynchus masou)*, and Amur catfish (*Silurus asotus*) (*P* < 0.05, Table 1 for statistically significant taxa, Table S5 for all taxa).

**Table 1.**
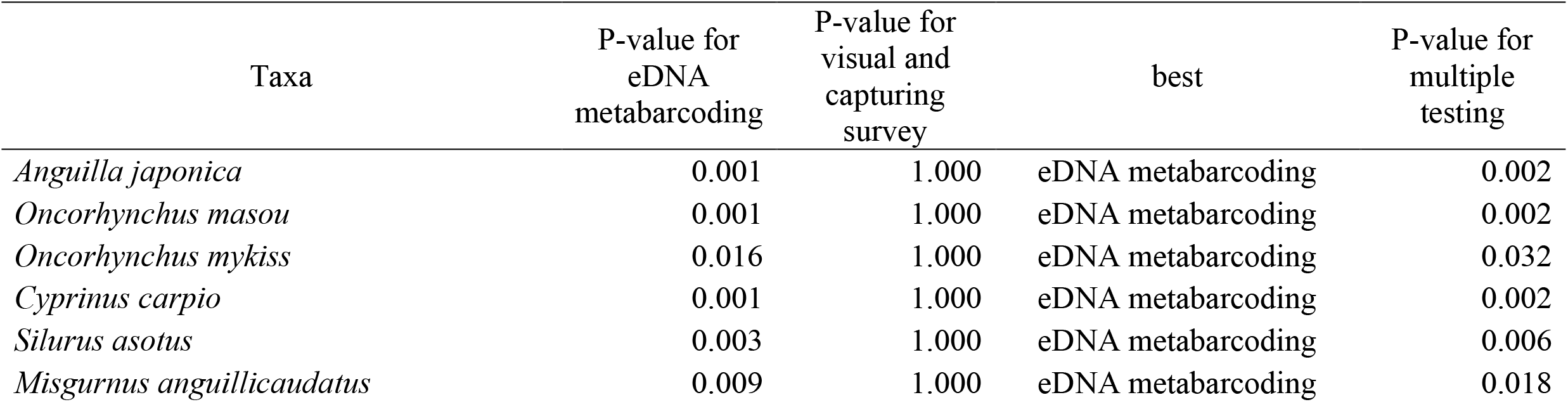
Indicator taxa analysis for the taxa which had significantly different detection frequencies between eDNA metabarcoding and visual/capture methods (P < 0.05). Best means preferred methods. P-values was calculated with 999 permutations after Sidak’s correction of the multiple testing.

### Nestedness and species turnover

We evaluated nestedness and turnover between communities as assessed using eDNA compared with visual/capture surveys using pairwise standardized effect size (pSES) calculated for all sites. Nestedness pSES was significantly positive without overlapping the zero-pSES 95% confidence interval (Fig. 3). The significantly negative pSES in species turnover indicated that taxa were not replaced in communities as assessed by eDNA compared to visual/capture surveys any more than would be expected to occur with random sampling.

**Figure 3.**
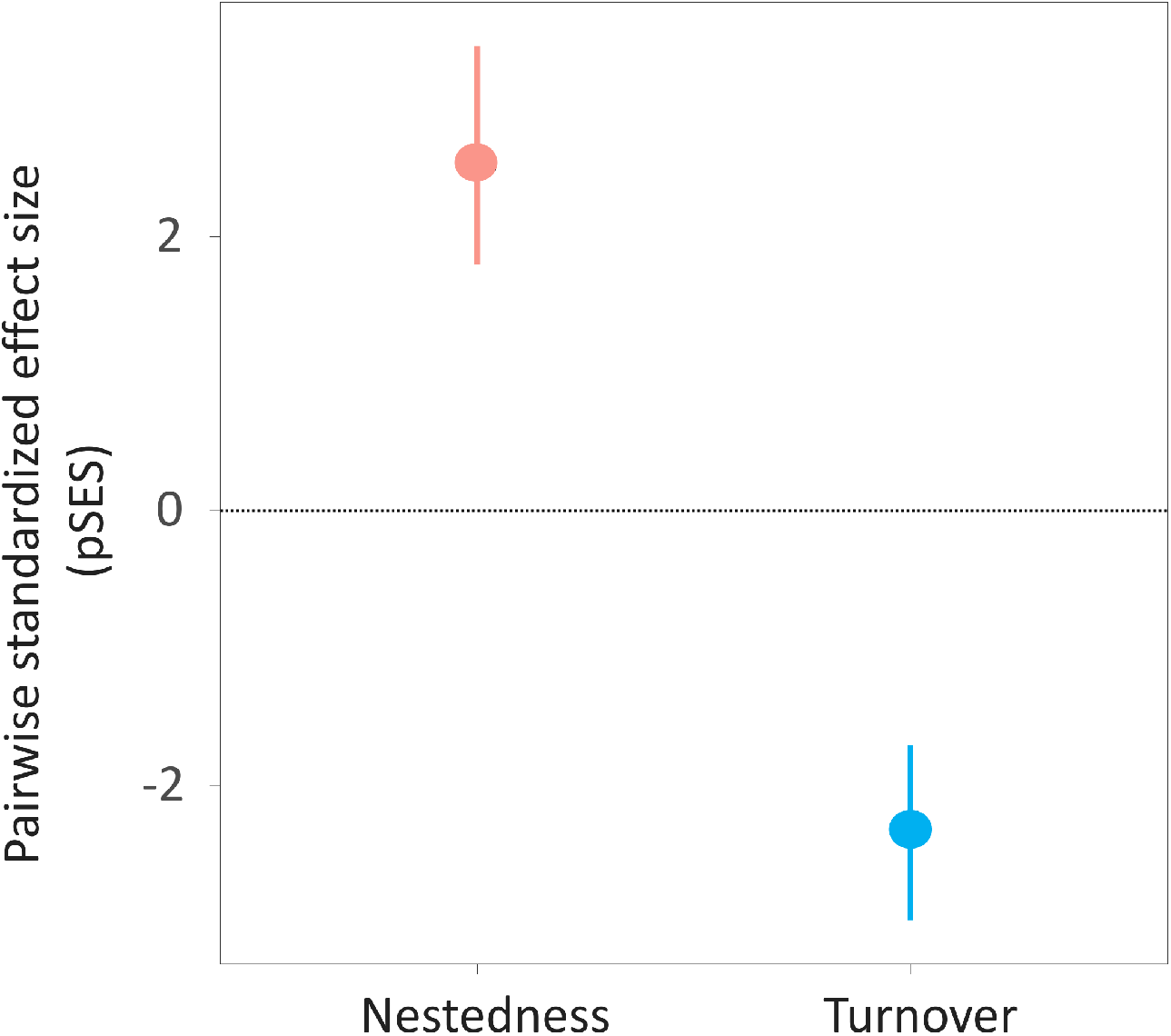
Species nestedness and turnover between the communities detected by eDNA metabarcoding and visual/capture methods, expressed as pairwise standardized effect sizes (SES). Error bars indicate 95% confidence intervals. The horizontal line represents SES=0, indicating a non-significant effect.

We compared the longitudinal nested structure and turnover strength (upstream to downstream) of the fish communities as assessed by eDNA metabarcoding and visual/capture surveys (Fig. 4). The longitudinal nested structure was not significantly different between both methods. The pSES values were significantly positive for both eDNA metabarcoding and visual/capture survey, indicating significant nested structures between segments, regardless of sampling method. Moreover, the pSES values were not significantly different between the eDNA metabarcoding and visual/capture survey, so the strength of the nested structures did not differ between the two methods (GLMM, *P* = 0.302). The species per segment were nested as downstream > mid-stream > upstream according to both methods (Fig. S5 for all species and S6 for each river, NOFD, *P* < 0.001). Longitudinal species turnover pSES values were not different between sampling methods (GLMM, p=0.28) and were significantly negative (Fig. 4b) indicating a lack of turnover whether communities were sampled by eDNA or visual/capture survey.

**Figure 4.**
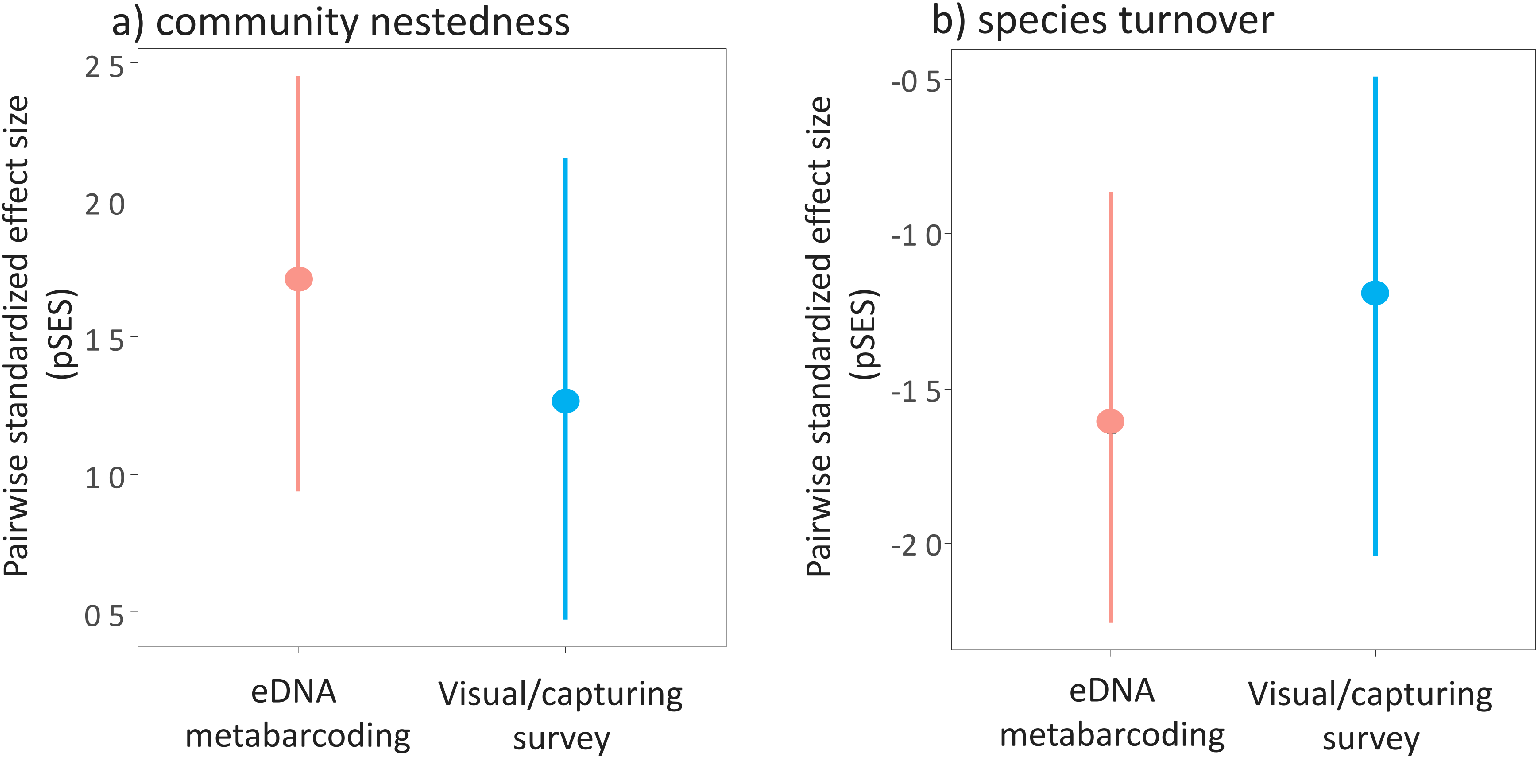
Effect sizes for (a) community nestedness and (b) species turnover among river segments as assessed by eDNA metabarcoding and visual/capture methods. The error bars indicate 95% confidence intervals

## Discussion

We here examined the nestedness and species turnover patterns of fish communities evaluated by eDNA metabarcoding in five rivers to evaluate the performance of eDNA metabarcoding. Our results suggested that the nestedness and species turnover from upstream to downstream did not significantly differ within a river. Therefore, we found that the both methods show similar levels of community nestedness and turnover within the rivers. In general, nestedness of species assemblages occurs when the communities obtained by the method estimating lower number of species are subsets of the communities estimated by other methods with higher species richness (Baselga 2010). Furthermore, the species turnover in the communities was very weak between the both methods.

eDNA metabarcoding has been reported to perform better than traditional methods in evaluating species richness (Deiner et al., 2016; Sato et al., 2017; Bylemans et al., 2018; Nakagawa et al., 2018; Yamamoto et al., 2018; Fujii et al., 2019). Nakagawa et al. (2018) investigated freshwater fish communities in 100 rivers and confirmed that the communities detected by eDNA metabarcoding were similar to the species lists observed in government-authorized monitoring. Furthermore, several eDNA metabarcoding studies on fish communities have been performed in other river systems (Bylemans et al., 2018), marine habitats (Yamamoto et al., 2018), and freshwater lakes (Sato et al., 2017; Fujii et al., 2019). Deiner et al. (2016) showed that river eDNA metabarcoding can reflect the regional community in a watershed, indicating that eDNA metabarcoding has high performance for gamma diversity evaluation. These studies indicated the great potential of eDNA metabarcoding as a useful tool for alpha and gamma diversity assessment by simply comparing the communities detected by eDNA metabarcoding and traditional surveys. Here, we support the previous literature (e.g., Nakagawa et al. 2018, Maechler et al. 2019) by showing patterns similar to those observed in alpha and gamma diversity in river macroinvertebrates. However, previous studies, excluding Maechler et al. (2019), did not evaluate performance in terms of nestedness and species turnover between eDNA metabarcoding and other community data. We also estimated nestedness, species turnover, and the capture bias of communities detected by eDNA metabarcoding and traditional methods in segment scale. We found that communities detected by traditional methods were nested within those detected by eDNA metabarcoding in a given river locale, with scarce species turnover between methods.

We especially focused on the variation of the community assemblages evaluated by eDNA metabarcoding compared to visual/capture survey. The two survey methods showed different sources of community variation. That is, eDNA metabarcoding detected significant differences between the rivers, while visual/capture survey detected significant differences between the segments rather than between the rivers. These differences may lead us to interpret the variation in community assemblages using the results from both survey methods. We recommend using eDNA for screening surveys because the method performs similarly to visual/capture surveys for estimating alpha and gamma diversity, although eDNA may fail to detect some species that can be captured or observed using traditional methods.

Using Jaccard index, a higher variation of the river fish communities was statistically detected by eDNA metabarcoding than by visual/capture survey and the variation of the community assemblages between segments could be significantly detected by visual/capture surveys but not by eDNA metabarcoding. While the results of Raup-Crick index, considering the alpha diversity of sites, showed the same results of the both methods. We further compared the indicator taxa for the communities obtained from both eDNA metabarcoding and visual/capture survey and concluded that several taxa, including eel, salmon, and catfish, were significantly better detected by eDNA metabarcoding, whereas non-indicator taxa were detected by visual/capture surveys. These results indicated that eDNA metabarcoding had higher detection frequency than visual/capture surveys in fish taxa detection. The community structures estimated by eDNA metabarcoding and visual/capture survey were slightly different, as reported in previous studies (e.g., Sato et al., 2017; Fujii et al., 2019), probably because of the differences in taxa-detection performances.

Discriminated taxa in this analysis included eel, salmon, and catfish, which mostly had larger body size and lower abundances in these rivers (Kawanabe, 2001; Nakabo, 2013), although we here could not separately examine the detected capture bias and actual pattern differences in species detection. In fact, the Japanese eel *Anguilla japonica*, was difficult to find by visual observation, probably due to its hiding behavior (Itakura et al., 2019). Such endangered species would be important as top predators (Nakabo, 2013). eDNA metabarcoding can evaluate the distribution of such rare and important taxa in fish communities better than traditional surveys. While we did not detect any indicator taxa by visual/capture surveys, two species *Lepomis macrochirus* and *Biwia zezera* were only detected by this method*. Lepomis macrochirus* often inhabit lentic systems and are rarely observed in rivers (Nakabo, 2013) and *Biwia zezera* are not widely distributed in this region (Hosokawa et al., 2007). In fact, we only observed *L. macrochirus* and *B. zezera* in downstream segments of the Kyuragi and Oze river, respectively.

We sampled eDNA from only two habitats at each study locale and examined the performances of eDNA metabarcoding. Thus, our understanding of some aspects of the fish community spatial structure in the rivers and the performance of community evaluation in local habitats, such as backwater, was still limited. In fact, Bylemans et al. (2018) found that river morphology in these habitats influenced the optimal sampling strategy for eDNA metabarcoding. Moreover, in backwater lakes, the performance of eDNA metabarcoding varied with different lake morphologies (Fujii et al. 2019). However, testing the usefulness of the eDNA metabarcoding for assessing river fish community biodiversity has been limited. Further research is needed to evaluate fish community spatial structure in rivers. In addition, we should consider whether eDNA recovered from a water sample came from an individual in the survey area. Previous studies have suggested the eDNA came from upstream (Deiner, K., & Altermatt 2014; Deiner et al. 2016). Therefore, the comparisons between eDNA metabarcoding and visual/capture methods could be using community data with different spatial scales. For example, visual/capture methods obtained community data from a 100-m reach but eDNA potentially detected the community in larger area than surveyed. However, the visual/capture survey also does not cover all species due to limited sampling time and efforts.

Community assessment with eDNA metabarcoding required much less effort in the field and detected the community in broader area than visual/capture survey. Therefore, eDNA metabarcoding may be a suitable method, especially for regional community patterns. Biodiversity testing using statistical frameworks, especially community nestedness and turnover, provided the quantitative evidence to compare the performance of eDNA metabarcoding and traditional surveys. eDNA methods for biodiversity assessment may provide more information to us, as shown here, but we should also pay attention to the unknown characteristics of eDNA, such as the origins, degradation, and transport of eDNA in water, which are still unknown in various habitats (Barnes & Turner, 2016; Seymour et al., 2018).

## Acknowledgments

MiSeq sequencing was conducted in the Department of Environmental Solution Technology, Faculty of Science and Technology, Ryukoku University, and we thank to H. Yamanaka and H. Sato for supporting the experiments involving MiSeq sequencing. Fish visual/capture surveys were permitted by the prefectures (No. 38 for Fukuoka, No. 3024 for Yamaguchi prefecture) for the survey period; 23 Sep. to 30 Nov. 2016. We conducted the fish survey under the guidance of Fisheries Adjustment Rules of the prefectures and Yamaguchi University. This study was supported by the Environment Research and Technology Development Fund (4-1602) of Environmental Restoration and Conservation Agency, Japan and JST-CREST (JPMJCR13A2).

## Data availability

All data of the MiSeq sequencing was shared in DRA (Accession number: DRA008090). Data was available in the supplementary material Table S1 and S2.

## Author contributions

HD, RI, and YA designed the study, RI, MG, TK, and YA contributed to field survey and sampling. SM, HD, RI, and MG contributed to molecular experiments. SM and HD analyzed the data and interpreted the results. HD, SM, and RI wrote the initial draft of the manuscript. All other authors critically reviewed the manuscript.

