## Supplemental Materials for "Estimation of biodiversity metrics by environmental DNA metabarcoding compared with visual and capture surveys of river fish communities"

Table S1 Taxa list evaluated by visual/capturing survey (v/c) and eDNA metabarcoding (eDNA). Excel file (SEM-Table S1).

Table S2 Miseq sequence reads for the samples. Excel file (SEM-Table S2).

Table S3 PERMANOVA results by Jaccard and Raup-Crick dissimilarity indices for community structures in the both methods, eDNA metabarcoding and visual/capturing survey, segments and rivers.

Table S4 PERMANOVA results by Jaccard and Raup-Crick dissimilarity indices for community structures in study rivers and sites.

Table S5 Indicator taxa analysis to determine which taxa had significantly different frequency between the both methods for all study taxa. Best means preferred methods. P-values was calculated with 999 permutations after Sidak's correction of the multiple testing.

Figure S1 Map for the five study rivers and the sampling sites in the study rivers.

Figure S2 The airborne photo for an example sampling.

Figure S3 The relationship between number of sequence reads and OUT numbers, i.e., rarefaction curves for the samples.

Figure S4 Nonmetric multidimensional scaling (NMDS) ordination with Raup-Crick index). Fish communities evaluated by the study rivers (shape) and each segment (colored) of the river. MDS stress was 0.2023.

Figure S5 Nestedness analysis among the samples in fish community.

Figure S6 Nestedness analysis among the samples in fish community in each river.

Table S3 PERMANOVA results by Jaccard and Raup-Crick dissimilarity indices for community structures in the both methods, eDNA metabarcoding and visual/capturing survey, segments and rivers.

a) PERMANOVA results by Jaccard dissimilarity indices

| Parameters | Df | Sums of Sqs | Mean Sqs | *F* | $R^2$ | *P* |
|---|---|---|---|---|---|---|
| Survey methods | 1 | 0.767 | 0.766 | 3.395 | 0.108 | 0.003 |
| Segments | 2 | 1.332 | 0.666 | 3.123 | 0.188 | 0.001 |
| Rivers | 4 | 2.068 | 0.517 | 2.574 | 0.292 | 0.001 |

b) PERMANOVA results by Raup-Crick dissimilarity indices

| Parameters | Df | Sums of Sqs | Mean Sqs | *F* | $R^2$ | *P* |
|---|---|---|---|---|---|---|
| Survey methods | 1 | 0.691 | 0.691 | 6.30 | 0.183 | 0.012 |
| Segments | 2 | 0.786 | 0.393 | 3.56 | 0.208 | 0.029 |
| Rivers | 4 | 2.697 | 0.674 | 15.8 | 0.717 | 0.001 |

Table S4 PERMANOVA results by Jaccard and Raup-Crick dissimilarity indices for community structures in segments and rivers for each survey method.

a) PERMANOVA results by Jaccard dissimilarity indices

| Methods | Parameters | Df | Sums of Sqs | Mean Sqs | *F* | $R^2$ | *P* |
|---|---|---|---|---|---|---|---|
| Visual/capturing survey | Segments | 2 | 1.075 | 0.537 | 2.606 | 0.303 | 0.011 |
| | Rivers | 4 | 1.271 | 0.318 | 1.394 | 0.358 | 0.120 |
| eDNA metabarcoding | Segments | 2 | 0.582 | 0.291 | 1.595 | 0.21 | 0.061 |
| | Rivers | 4 | 1.245 | 0.311 | 2.039 | 0.449 | 0.001 |

b) PERMANOVA results by Raup-Crick dissimilarity indices

| Methods | Parameters | Df | Sums of Sqs | Mean Sqs | *F* | $R^2$ | *P* |
|---|---|---|---|---|---|---|---|
| Visual/capturing survey | Segments | 2 | 0.284 | 0.142 | 0.863 | 0.126 | 0.506 |
| | Rivers | 4 | 1.493 | 0.373 | 4.87 | 0.661 | 0.018 |
| eDNA metabarcoding | Segments | 2 | 0.400 | 0.200 | 1.989 | 0.249 | 0.214 |
| | Rivers | 4 | 1.401 | 0.350 | 16.81 | 0.870 | 0.002 |

Table S5 Indicator taxa analysis to determine which taxa had significantly different frequency between the both methods for all study taxa. Best means preferred methods. P-values was calculated with 999 permutations after Sidak's correction of the multiple testing. The bold letters indicated the significant species in the p-value for multiple testing.

| Taxa | P-value for eDNA metabarcoding | P-value for visual and capturing survey | best | P-value for multiple testing |
|---|---|---|---|---|
| *Anguilla japonica* | **0.001** | **1.000** | **eDNA metabarcoding** | **0.002** |
| *Plecoglossus altivelis altivelis* | 0.146 | 0.963 | eDNA metabarcoding | 0.271 |
| *Oncorhynchus keta* | 0.516 | 1.000 | eDNA metabarcoding | 0.766 |
| *Oncorhynchus masou* | **0.001** | **1.000** | **eDNA metabarcoding** | **0.002** |
| *Salvelinus leucomaenis* | 0.218 | 1.000 | eDNA metabarcoding | 0.388 |
| *Oncorhynchus mykiss* | **0.016** | **1.000** | **eDNA metabarcoding** | **0.032** |
| *Oryzias latipes* | 0.509 | 0.826 | eDNA metabarcoding | 0.759 |
| *Tanakia limbata* | 0.493 | 0.892 | eDNA metabarcoding | 0.743 |
| *Squalidus gracilis gracilis* | 0.376 | 0.866 | eDNA metabarcoding | 0.611 |
| *Tribolodon hakonensis* | 0.075 | 0.987 | eDNA metabarcoding | 0.144 |
| *Opsariichthys platypus* | 0.323 | 0.910 | eDNA metabarcoding | 0.542 |
| *Rhodeus smithii smithii* | 0.771 | 0.753 | visual/capturing | 0.939 |
| *Acheilognathus rhombeus* | 0.239 | 1.000 | eDNA metabarcoding | 0.421 |
| *Pseudogobio esocinus esocinus* | 0.798 | 0.485 | visual/capturing | 0.735 |
| *Sarcocheilichthys variegatus variegatus* | 0.764 | 0.750 | visual/capturing | 0.938 |
| *Candidia temminckii* | 0.515 | 1.000 | eDNA metabarcoding | 0.765 |
| *Carassius* sp. | 0.029 | 0.992 | eDNA metabarcoding | 0.057 |
| *Carassius cuvieri* | 0.114 | 1.000 | eDNA metabarcoding | 0.215 |
| *Cyprinus carpio* | **0.001** | **1.000** | **eDNA metabarcoding** | **0.002** |
| *Squalidus chankaensis tsuchigae* | 0.513 | 1.000 | eDNA metabarcoding | 0.763 |
| *Hemibarbus longirostris* | 0.491 | 0.881 | eDNA metabarcoding | 0.741 |
| *Biwia zezera* | 1.000 | 0.490 | visual/capturing | 0.740 |
| *Phoxinus oxycephalus jouyi* | 0.128 | 0.983 | eDNA metabarcoding | 0.240 |
| *Gnathopogon* sp. | 0.305 | 0.944 | eDNA metabarcoding | 0.517 |
| *Hemibarbus* sp. | 0.188 | 0.957 | eDNA metabarcoding | 0.341 |
| *Rhodeus ocellatus kurumeus* | 0.771 | 0.753 | visual/capturing | 0.939 |
| *Opsariichthys uncirostris uncirostris* | 0.513 | 1.000 | eDNA metabarcoding | 0.763 |
| *Pungtungia herzi* | 0.495 | 0.777 | eDNA metabarcoding | 0.745 |
| *Pseudorasbora parva* | **0.017** | **1.000** | **eDNA metabarcoding** | **0.034** |
| *Tanakia lanceolata* | 0.244 | 1.000 | eDNA metabarcoding | 0.428 |

| | | | | |
|---|---|---|---|---|
| *Liobagrus reini* | 0.817 | 0.520 | visual/capturing | 0.770 |
| *Tachysurus nudiceps* | 0.678 | 0.631 | visual/capturing | 0.864 |
| *Tachysurus aurantiacus* | 0.677 | 0.673 | visual/capturing | 0.893 |
| *Silurus asotus* | **0.003** | **1.000** | **eDNA metabarcoding** | **0.006** |
| *Misgurnus anguillicaudatus* | **0.009** | **1.000** | **eDNA metabarcoding** | **0.018** |
| *Cobitis takatsuensis* | 0.494 | 1.000 | eDNA metabarcoding | 0.744 |
| *Cobitis biwae* | 0.689 | 0.678 | visual/capturing | 0.896 |
| *Cobitis striata* | 0.513 | 1.000 | eDNA metabarcoding | 0.763 |
| *Cobitis matsubarae* | 0.483 | 0.830 | eDNA metabarcoding | 0.733 |
| *Mugil cephalus cephalus* | 0.516 | 1.000 | eDNA metabarcoding | 0.766 |
| *Chelon haematocheilus* | 0.516 | 1.000 | eDNA metabarcoding | 0.766 |
| *Cottus pollux* | 0.501 | 0.777 | eDNA metabarcoding | 0.751 |
| *Cottus kazika* | 0.705 | 0.698 | visual/capturing | 0.909 |
| *Micropterus salmoides* | 0.504 | 1.000 | eDNA metabarcoding | 0.754 |
| *Coreoperca kawamebari* | 0.685 | 0.635 | visual/capturing | 0.867 |
| *Lateolabrax japonicus* | 0.253 | 1.000 | eDNA metabarcoding | 0.442 |
| *Lepomis macrochirus* | 1.000 | 0.450 | visual/capturing | 0.698 |
| *Channa argus* | 0.304 | 0.950 | eDNA metabarcoding | 0.516 |
| *Odontobutis* sp. | 0.303 | 0.917 | eDNA metabarcoding | 0.514 |
| *Gymnogobius urotaenia* | 0.483 | 0.890 | eDNA metabarcoding | 0.733 |
| *Gymnogobius petschiliensis* | 0.516 | 1.000 | eDNA metabarcoding | 0.766 |
| *Rhinogobius* sp. | 0.092 | 0.988 | eDNA metabarcoding | 0.176 |
| *Rhinogobius flumineus* | 0.994 | 0.059 | visual/capturing | 0.115 |
| *Rhinogobius similis* | 0.662 | 0.642 | visual/capturing | 0.872 |
| *Tridentiger* sp. | 0.359 | 0.870 | eDNA metabarcoding | 0.589 |

Note: The p values of *Biwia zezera* and *Lepomis macrochinus* were not calculated because of their low presence (one detection per 30 samples.).

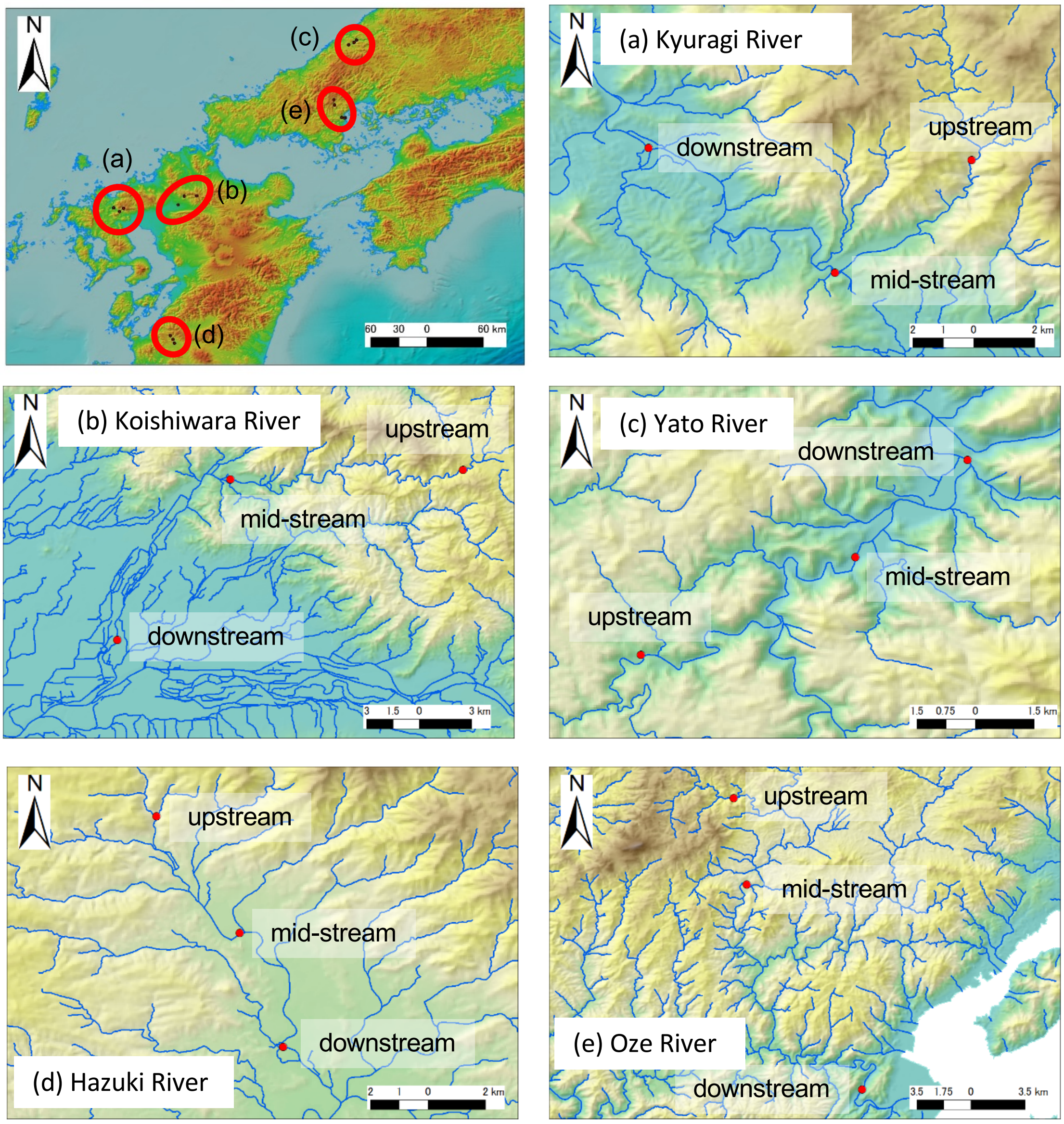


Figure S1 Map for the five study rivers and the sampling sites in the study rivers.

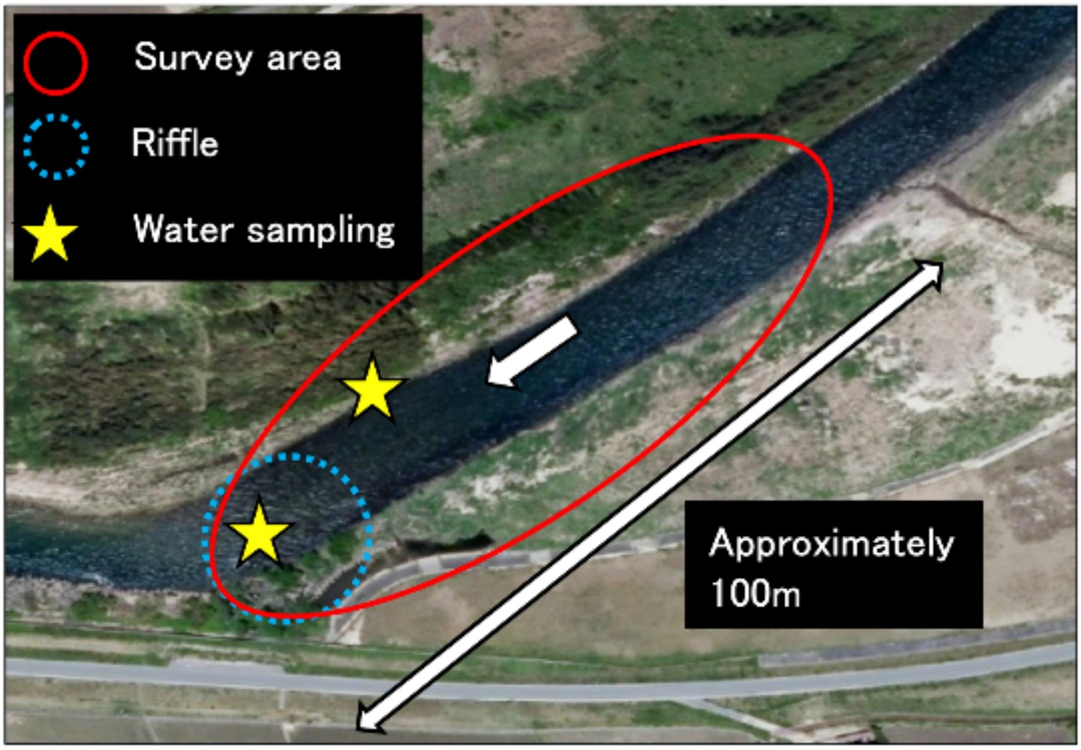


Figure S2 The airborne photo for an image of sampling site (in Saba river, Japan).

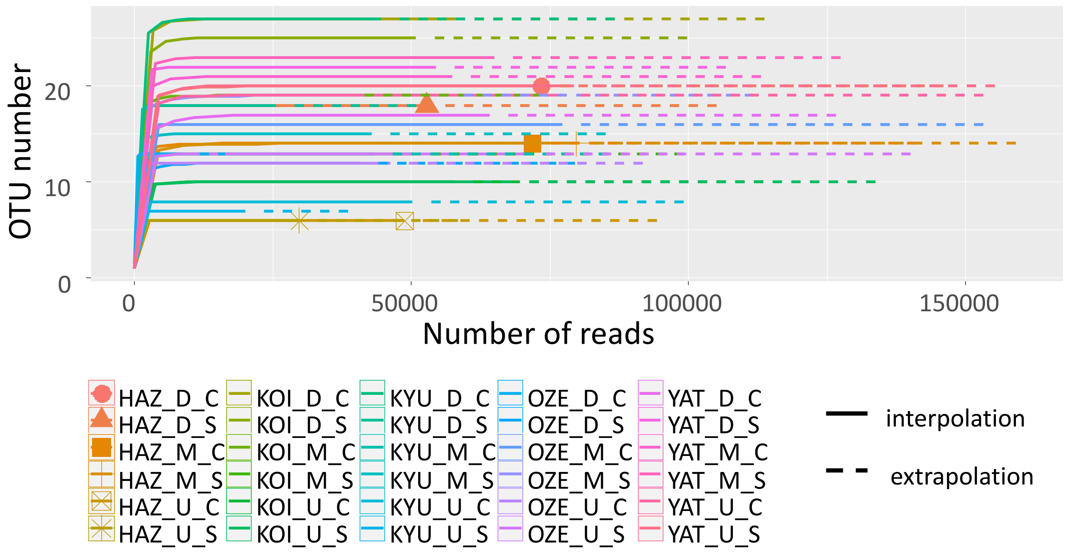


Figure S3 The relationship between number of sequence reads and OUT numbers, i.e., rarefunction curves for the samples. In the legend, the first three characters in the top names indicated the river names (KOI= Koi river, KYU=Kyuragi river, HAZ= Hazuki river, OZE=Oze river, YAT=Yato river), and D, M, U after the river names indicated downstream, mid-stream, and upstream, and the last letters (M and T) indicated eDNA metabarcoding and traditional survey (visual/capturing), respectively, e.g., KOI_D_M means KOI river-downstream-eDNA metabarcoding

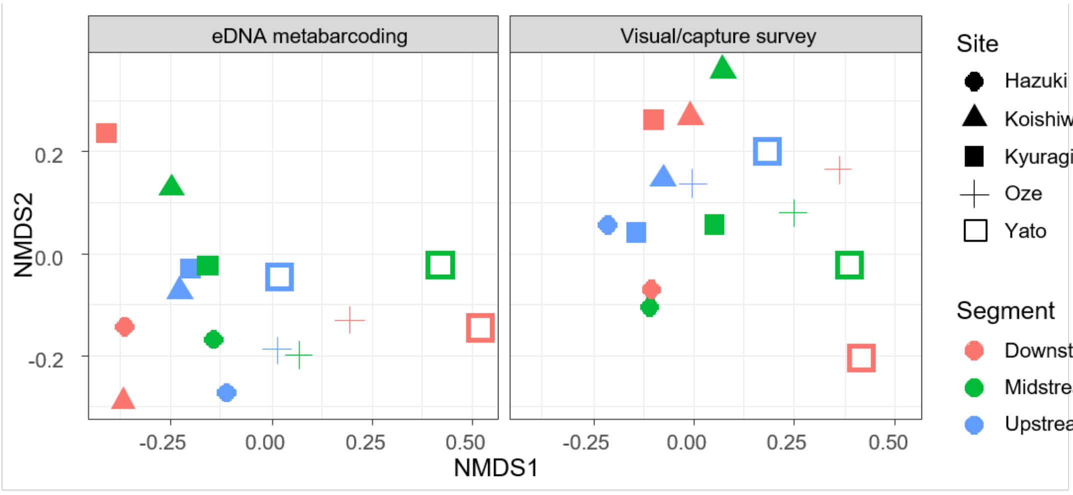


Figure S4 Nonmetric multidimensional scaling (NMDS) ordination with Raup-Crick index). Fish communities evaluated by the study rivers (shape) and each segment (colored) of the river. MDS stress was 0.2023.

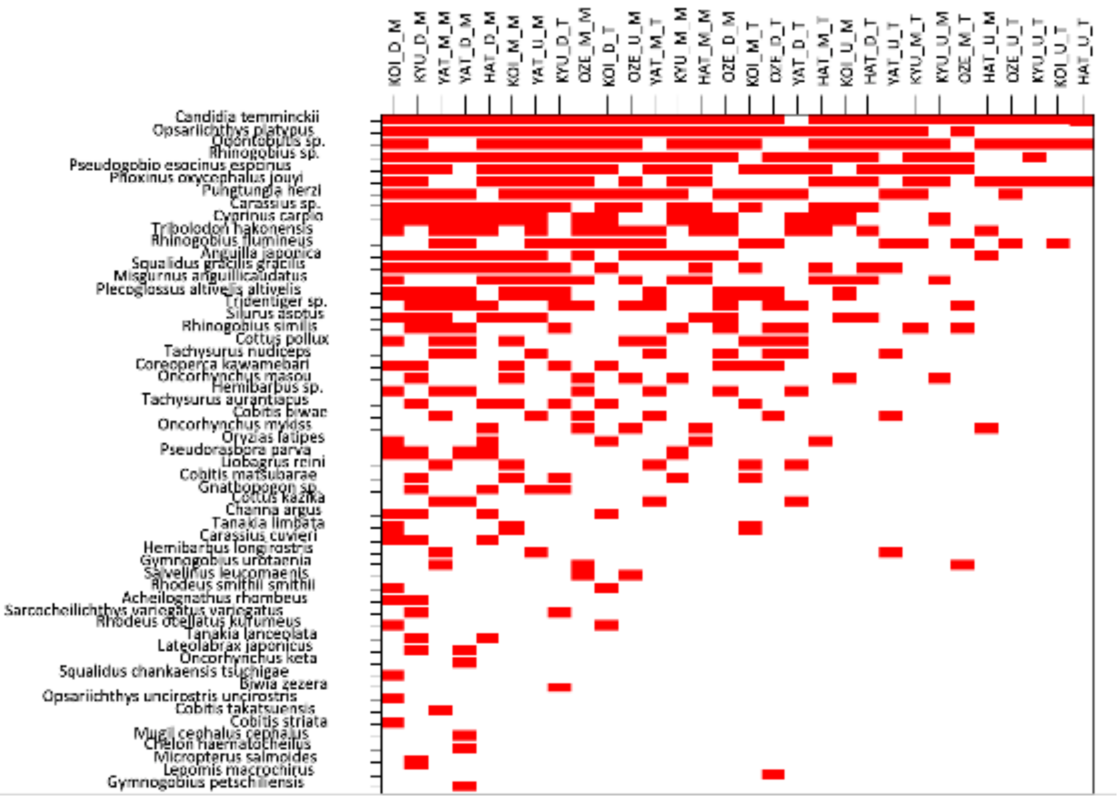


Figure S5 Nestedness analysis among the samples in fish community. The first three characters in the top names indicated the river names (KOI= Koi river, KYU=Kyuragi river, HAT= Hazuki river, OZE=Oze river, YAT=Yato river), and D, M, U after the river names indicated downstream, mid-stream, and upstream, and the last letters (M and T) indicated eDNA metabarcoding and traditional survey (visual/capturing), respectively, e.g., KOI_D_M means KOI river-downstream-eDNA metabarcoding

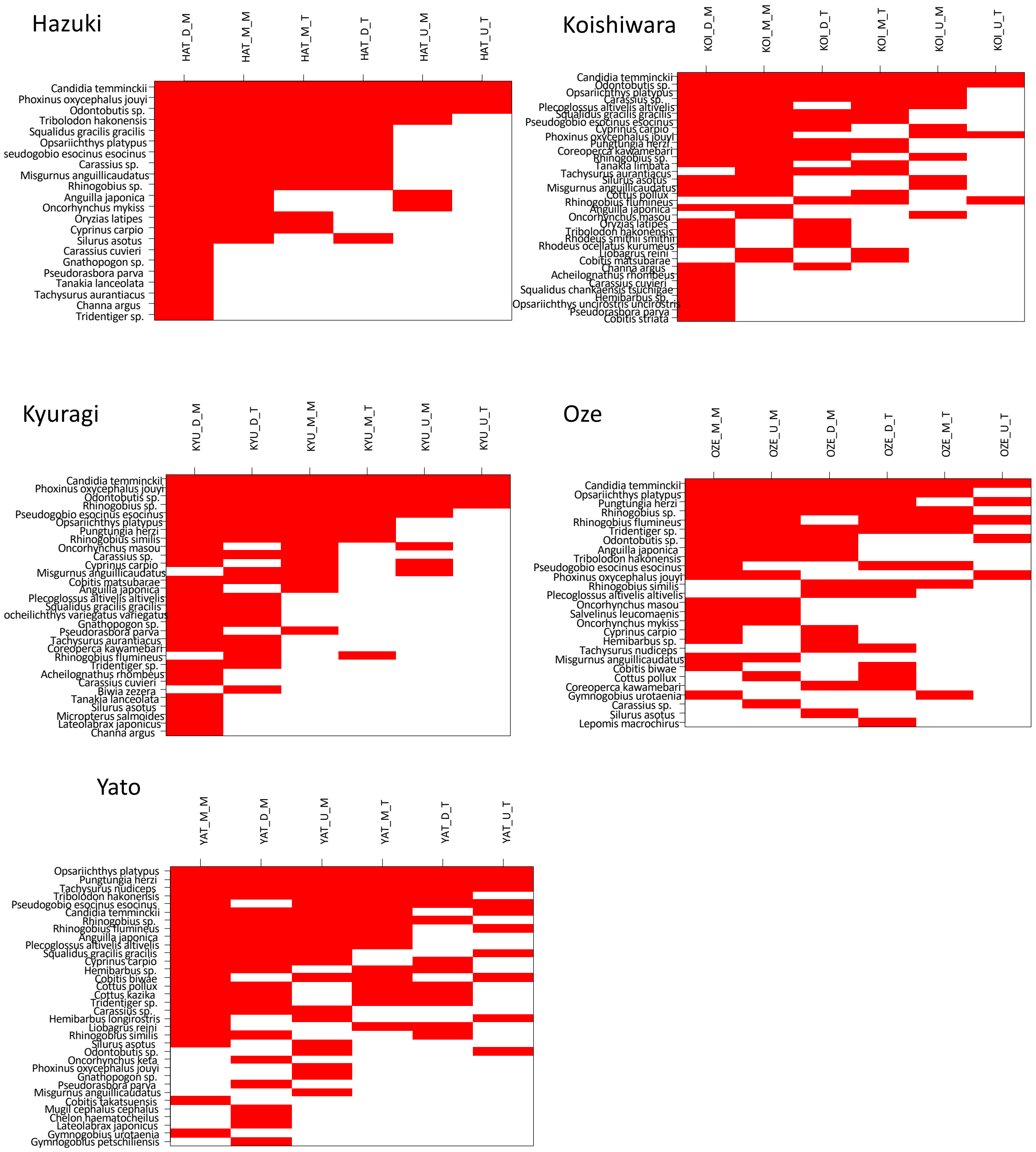


Figure S6 Nestedness analysis among the samples in fish community in each river. The first three characters in the top names indicated the river names, and D, M, U after the river names indicated downstream, mid-stream, and upstream, and the last letters (M and T) indicated eDNA metabarcoding and traditional survey (visual/capturing), respectively, e.g., KOI_D_M means KOI river-downstream-eDNA metabarcoding
